# Identifying female phenotypes that promote behavioral isolation in a sexually dimorphic species of fish (*Etheostoma zonale*)

**DOI:** 10.1101/2020.04.20.051714

**Authors:** Natalie S. Roberts, Tamra C. Mendelson

## Abstract

In sexually dimorphic species characterized by exaggerated male ornamentation, behavioral isolation is often attributed to female preferences for conspecific male signals. Yet, in a number of sexually dimorphic species, male mate choice also results in behavioral isolation. In many of these cases, the female traits that mediate species boundaries are unclear. Females in sexually dimorphic species typically lack many of the elaborate traits that are present in males and that are often diagnostic of species. In a diverse and largely sexually dimorphic group of fishes called darters (Percidae: *Etheostoma*), male mate choice contributes to behavioral isolation between a number of species; however, it is not clear which female traits males prefer. In the current study, we identified the dominant female pattern for two sympatric species, *Etheostoma zonale* and *E. barrenense*, using pattern energy analysis, and we used discriminate function analysis to identify which aspects of female patterning can reliably classify species. We then tested the role of female features in male mate choice for *E. zonale*, by measuring male preference for computer animations displaying the identified (species-specific) conspecific features as well as the dominant male pattern that is preferred by females. We found that the region above the lateral line is important in mediating male mate preferences, with males spending significantly more time with animations exhibiting conspecific female patterning in this region than with animations exhibiting heterospecific female patterning. Our results suggest that the aspects of female phenotypes that are the target of male mate choice are different from the male phenotypes that characterize species. This research highlights the importance of using objective measures in the study of behavioral isolation via male mate choice.

## Introduction

Behavioral isolation is the reduction of gene flow between populations or species due to differences in courtship signals and preferences, and it is often considered one of the most important barriers to gene flow between closely related animal species (Coyne & Orr, 2004). Identifying the courtship signals whose differences contribute to behavioral isolation therefore is fundamental in understanding the dynamics of many species boundaries. In species characterized by extreme sexual dimorphism, behavioral isolation is most often attributed to female preferences for elaborate conspecific male traits (Kraaijeveld et al., 2011), as female preferences are thought to be primarily responsible for male trait elaboration (Andersson, 1994). Traits that promote behavioral isolation via female choice tend to be male ornaments that are diagnostic of species (e.g. red vs. blue coloration in haplochromine cichlids, Seehausen & van Alphen, 1998; pulse rate in *Gryllus* and *Laupala* crickets, Gray & Cade, 2000, Mendelson & Shaw, 2006; visual and vocal signals in *Passerina spp.*, Baker & Baker, 1990), which females generally lack or possess in reduced forms.

Behavioral isolation via male mate choice has been studied primarily in species that are not known for striking sexual dimorphism, such as *Drosophila* spp. and Lepidoptera (von Schilcher & Dow 1977; Roelofs & Camaeu 1969; Cobb & Jallon 1990; Coyne & Oyama 1995; Jiggins et al. 2004). In these lineages, the traits that males prefer are often present in both sexes and are generally under strong ecological selection (Jiggins, 2008; Merrill et al., 2012; Chung & Carroll, 2015). For example, wing color patterns that promote assortative mating in *Heliconius* butterflies are present in both males and females and also signal distastefulness to predators (Jiggins et al., 2004). Cuticular hydrocarbons that function in male preference for conspecifics in *Drosophila* spp. are sometimes sexually dimorphic and also help prevent against desiccation (Chung et al. 2014; Combs et al., 2018).

Fewer studies examine the female signals that mediate male preferences for conspecific mates in sexually dimorphic species, when females are drab and unornamented. It is understandably much harder to test which female traits prevent interbreeding if relevant female traits are difficult to identify. However, recent studies in the sexually dimorphic fish genus *Etheostoma* (darters) suggest that male preferences contribute as much as or more than female preferences to behavioral isolation among species in this group (Zhou et al., 2015; Roberts & Mendelson, 2017; Moran et al., 2017) and that male preferences may evolve earlier than female preferences in several allopatric species pairs (Mendelson et al., 2018). Preferences for conspecific mates are based largely on visual signals, with both females and males preferring conspecific stimuli when only visual cues are presented (Williams & Mendelson, 2010; Roberts & Mendelson, 2017). These findings suggest that females possess species-identifying visual features, even if they are phenotypically much less distinct than males, and that males prefer species-specific female phenotypes.

*Etheostoma zonale* and *E. barrenense* are sympatric species of darters that have distinct, species-specific male coloration and pattern; females of both species are unornamented by comparison. Male *E. zonale* have alternating green and yellow bars along the body while male *E. barrenense* display primarily red-orange coloration with black blotches fused along the lateral line to form a stripe (Fig 1a, 1b). Females display the patterning of conspecific males but only muted coloration, if any at all (Fig. 1c, 1d). In a previous study aimed to identify the visual cues that explain female preference for conspecific males in this species pair, Williams & Mendelson (2011) used hand-painted motorized models and showed that females of both species prefer conspecific male color (green versus red models) and pattern (bars versus stripe). In contrast, the phenotypes that drive male preference for conspecific females remain unclear.

**Figure 1.**
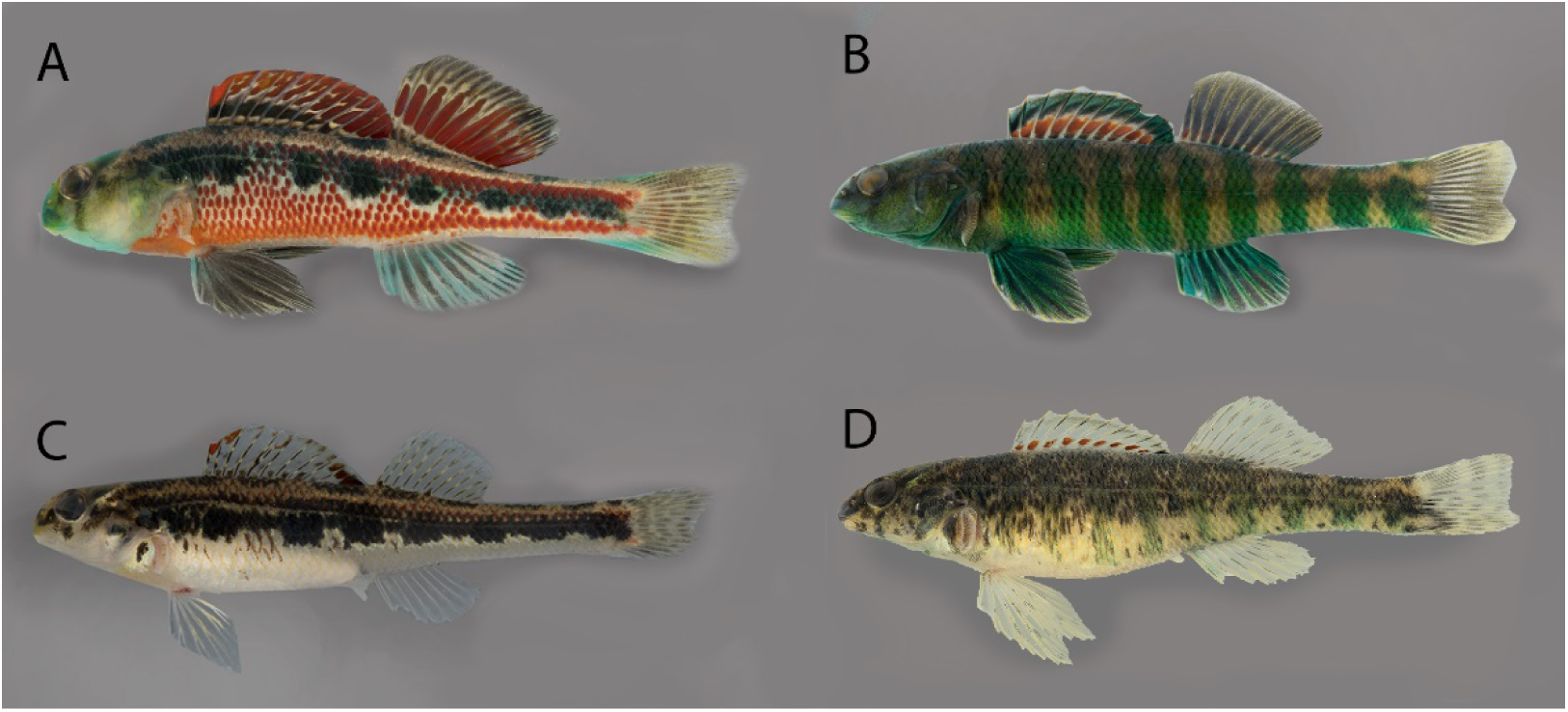
Representative photos taken during breeding season of male (A) and female (C) *E. barrenense*, and male (B) and female (D) *E. zonale*.

In the current study, we aimed to determine which aspects of female phenotype contribute to behavioral isolation via male preferences in this species pair. We first used pattern energy analysis to identify the dominant female pattern (Stoddard & Stevens, 2010; Troscianko & Stevens, 2015). Pattern energy analysis analyzes the contribution of different marking sizes to a given pattern and has been used to quantify various aspects of pattern attributes (e.g. marking size, coverage) in studies of host-parasite egg matching (Stoddard & Stevens, 2010). We then used discriminant function analysis to determine which features of the dominant female region identified via pattern energy analysis can reliably classify species. Female pattern features that could reliably classify species were then tested in behavioral assays to measure their effect on behavioral isolation between *E. zonale* and *E. barrenense*. We also identified male dominant pattern using pattern energy analysis and tested male preference for the identified male-specific dominant pattern applied to female *E. zonale*. We utilized 3D digital animations in behavioral assays, allowing us to test whether addition of the conspecific features to heterospecific animations would increase male preference.

## Methods

Animations were created and behavioral assays were conducted in 2018 and 2019. Experimental protocols changed slightly between the two years based on preliminary results obtained in 2018 and to improve overall testing procedures. All experimental methods outlined below apply to both 2018 and 2019 trials unless specifically noted as Year 1 (2018 trials) or Year 2 (2019 trials).

### Fish collection and care

Fish used for photography and pattern energy analysis were collected in 2017 between March – April. *Etheostoma zonale* were collected from the Middle Fork Red River, Powell Co., KY (37.815002, −83.71877) and Line Creek, Monroe Co., KY (36.651835, −85.820182), and *E. barrenense* were collected from the East Fork Barren River, Monroe Co., KY (36.745964, - 85.696728). Males used for behavioral assays were collected from Line Creek in Clay Co., TN (36.606639, −85.745970) and Monroe Co., KY (36.651835, −85.820182) on 22 March 2018 and from the same site in KY the following year on 23 March 2019. All fish were transported to the University of Maryland Baltimore County and housed in a recirculating aquarium system (Aquatic Habitats, Inc., Apopka, FL, USA) until euthanized for photography or used in behavioral assays. Water temperature, conductivity, and pH for fish housing approximated conditions in the natural habitat. Fish were maintained on a diet of live black worms provided once daily. Permission to collect fish was granted by the Kentucky Department of Fish and Wildlife Resources (2017: #SC1711121, 2018: #Sc1811149, 2019: #SC1911185) and the Tennessee Wildlife Resource Agency (Scientific Collection Permit #1424). Authority to work with live animals was granted by the IACUC (OLAW Assurance Number D16-00462, Protocol TM011061821).

### Quantifying focal regions: pattern energy analysis

RAW photographs of female (N = 38) and male (N = 27) *E. zonale* and female (N = 29) and male (N = 12) *E. barrenense* were taken with a macrophotography setup using a Canon Mark IV camera with a Canon EF 100mm f/2.8 L macro lens (Canon USA, Inc., Lake Success, NY, USA). Fish were positioned laterally and pinned with all fins extended (with the exception of the pectoral fin which was removed in order to provide an unobstructed view of body color and pattern) in 10% formalin for approximately 10 minutes. The camera was mounted on a Cognisys Stackshot Extended Macro Rail (Cognisys Inc., Traverse City, MI, USA) and set to focus the camera on the highest focal plane of each fish. Photographs were then taken at 0.5mm intervals until the lowest focal plane of the fish was in focus, generating a series of images at different focal planes. These images were then stacked using Zerene Stacker (Zerene Systems, Richland, WA, USA) to create a single high-resolution image for each fish and converted into darter color space using the Image Analysis and Calibration Toolbox plugin in ImageJ (Troscianko & Stevens, 2015; Roberts et al., 2017).

To identify the dominant marking pattern, we used the pattern energy analysis function in the ImageJ plugin (Troscianko & Stevens, 2015). Pattern energy analysis filters an original image into a series of other images, each representing a different spatial scale (Barbosa et al., 2008; Chiao et al., 2009; Stoddard & Stevens, 2010). This process is similar to a sieve, in which different filter sizes allow different aspects of an image to be retained, with the sum of all filtered images closely approximating the original image. Pattern energy is then calculated as the standard deviation of all pixel values for each filtered image, with the filter size that has the highest pattern energy (i.e. highest standard deviation of pixel values) being considered the dominant marking size (Stoddard & Stevens, 2010; Troscianko & Stevens, 2015). We filtered images using a 1.6 step multiplier, starting at a 4 pixel scale to generate 13 filtered images of each individual, with pattern energy calculated for each filtered image. We averaged pattern energy at each spatial scale across all individuals of each sex and species to identify the species-specific dominant marking filter size for males and females separately (i.e. the averaged filter size with the highest pattern energy for each sex and species; Fig. 2). In the filtered image corresponding to the sex- and species-level dominant marking size, we used the threshold tool in ImageJ to select the lowest pixel values (i.e. darker areas) until 25% of total pixel values were selected. We term this 25% selection the “focal region.”

**Figure 2.**
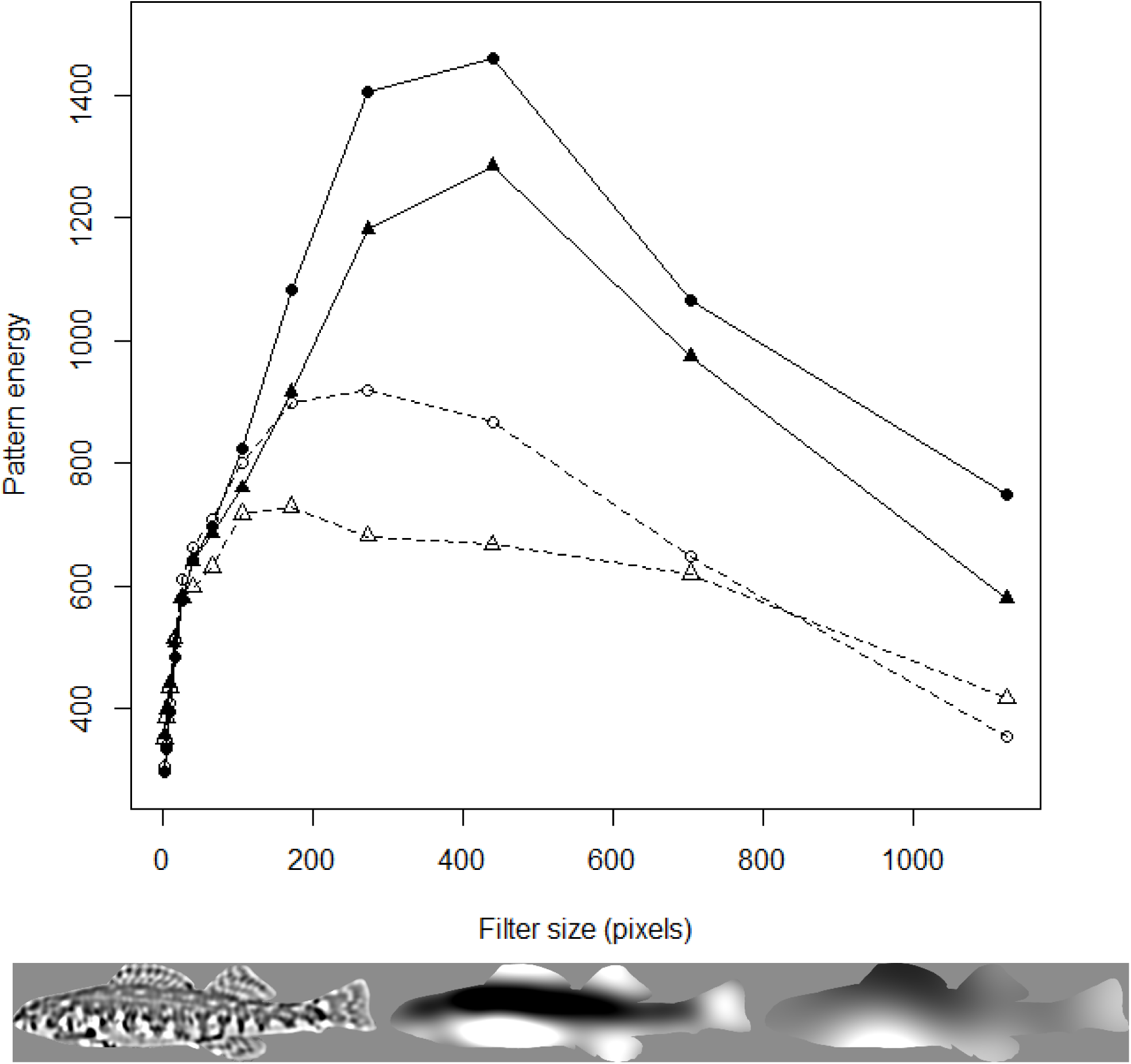
Average pattern energy spectra for *E. barrenense* (females: closed circles; males: open circles) and *E. zonale* (females: closed triangles; males: open triangles) across 13 filter sizes. Image series below shows an example of a female *E. zonale* photograph processed at different filter sizes (from left to right: 67 pixels, 440 pixels, and 1,126 pixels). The middle image (440 pixels) is the filter size with the highest pattern energy, indicating this is the dominant marking size.

The 25% cutoff value was somewhat arbitrarily chosen; however, using the 25% cutoff value for images of females appeared to capture nearly the entire region above the lateral line identified as a distinct marking in the filtered image with the highest pattern energy (Fig. 3a, 3c). For male *E. zonale* and *E. barrenense*, it captured the barred and stripe patterning, respectively, known to be preferred by females (Fig. 3b, 3c; Williams & Mendelson, 2011, 2013), suggesting that the 25% cutoff value can identify behaviorally relevant body patterns. Ideally, we would test a range of cutoff values in behavioral assays; however, the short (3 month) breeding season of darters is prohibitive. Thus, through pattern energy analysis and using a 25% threshold to select the dominant marking, we identified the female focal region as the flank above the lateral line for both *E. barrenense* and *E. zonale* (Fig. 3a, 3c).

**Figure 3.**
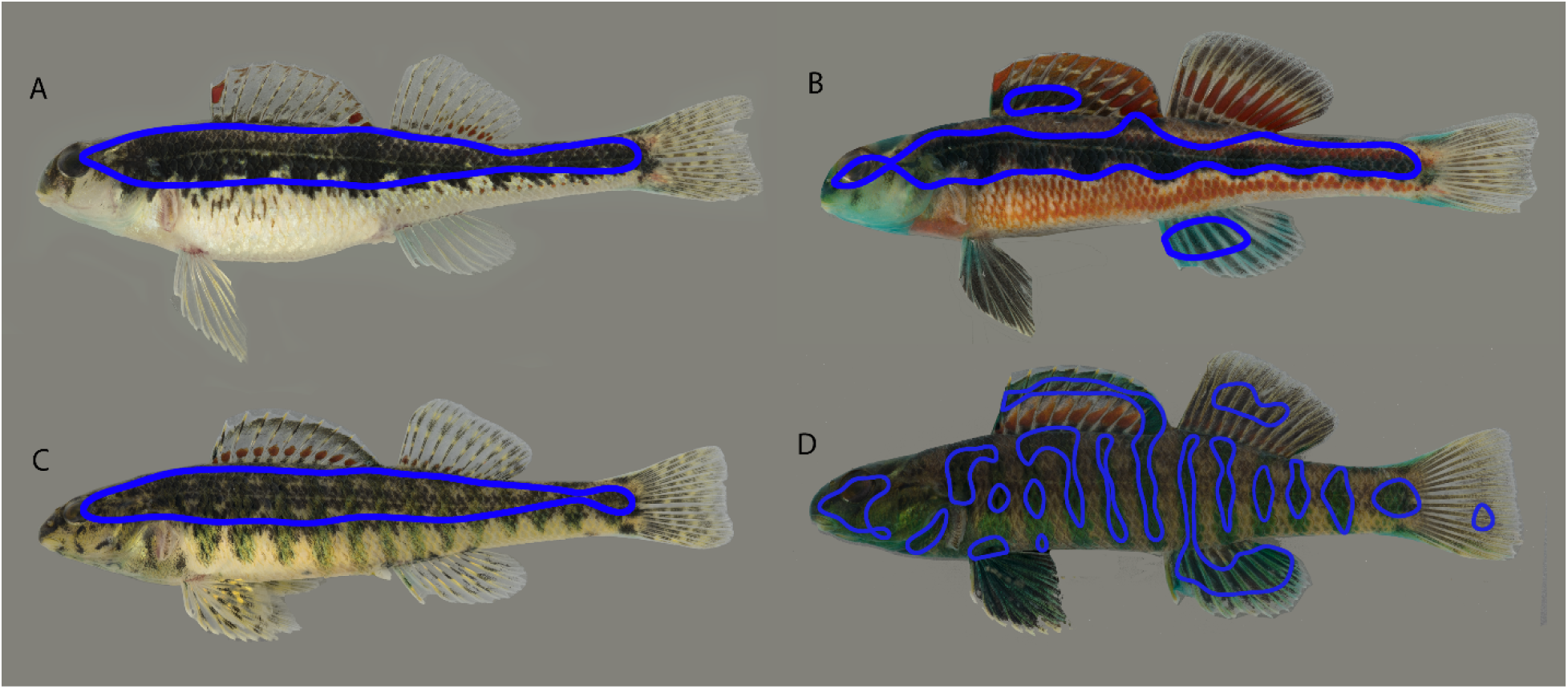
Representative images of focal regions (outlined in blue) for female (A) and male (B) *E. barrenense* and female (C) and male (D) *E. zonale* obtained by selecting 25% of the total body area representing the dominant pattern indicated for females by pattern energy analysis.

### Discriminant function analysis of female focal regions

We quantified the color, luminance, size, and shape of the *E. zonale*-specific female focal regions, in order to independently measure the effect of each of these focal region features in species classification (using discriminate function analysis) and male mate choice (using behavioral assays). To calculate focal region color and luminance, we estimated quantal catch from the darter medium wavelength-sensitive (MWS) and long wavelength-sensitive (LWS) cone types using the ImageJ plugin (Gumm et al., 2012; Roberts et al., 2017). Color was defined as the difference in quantal catch between the LWS and MWS cone types (Williams & Mendelson, 2013) and luminance was calculated as quantal catch from LWS types (Marshall & Jennings, 2003). Focal region size was quantified as the total number of pixels contained within the focal region. Focal region shape was quantified using elliptical Fourier analysis (Kuhl & Giardina, 1982; Rohlf & Archie, 1984; McLellan & Endler, 1998) which breaks down a shape outline into mathematically defined ellipses (i.e. harmonics) of different sizes, orientations, and eccentricities to describe increasingly finer shape details. This method has been used to classify complex biological shapes such as corals (*Pseudopterogorgia spp.*, Carlo et al., 2011) and face patterns in guenon monkeys (tribe: Cercopithecini, Allen & Higham, 2015). Using the program Shape (ver 1.3.; Iwata & Ukai, 2002), we extracted 20 harmonics, which achieved a close visual match to segmented shape, and then performed PCA to reduce shape descriptors.

To determine which of the independent features of the female focal region (i.e. color, luminance, size, or shape) could accurately predict species identity we used linear discriminant function analysis (DFA) in R (ver. 3.5.0; R Core Development Team 2015) using the package MASS and code written by R. Mundry (MPA for Evolutionary Anthropology, Leipzig, Germany). DFA attempts to classify groups on the basis of a set of predictor variables. We evaluated classification performance by evaluating whether our DFA model was able to correctly classify species significantly better than when compared to a null model. The null model is created by randomly assigning observations to either group (i.e. *E. zonale* or *E. barrenense*), irrespective of their actual group membership, and then performing DFA analysis on the randomized dataset. Thus, for the null model there should be no discriminability between groups. The classification accuracy of the observed data can then be compared to the null model to assess whether classification performance is significantly better than the null model. In addition to performing DFA on each measured focal region feature, we also performed DFA analysis on a model that included all focal region measurements, i.e., focal region color, luminance, size, and shape.

DFA analysis showed that for females, focal region color (78.9% correctly identified, *P* < 0.05), shape (89.5% correctly identified, *P* < 0.01), and the model with all focal region measures (91.5% correctly identified, *P* < 0.05) reliably classified species identity. We therefore tested the effect of these three features (color, shape, and an unaltered focal region) on male preferences in behavioral assays.

### Animations

Creation of 3D animations followed methods in Roberts et al. (2019). Three-dimensional digital models of *E. zonale* and *E. barrenense* were purchased from Turbosquid (product ID female *E. zonale*: 928023; female *E. barrenense*: 957636) and imported into the animation software program Autodesk Maya (Autodesk Inc, San Rafael, CA, USA). To create a path for models to follow, we imported video of live darters into Maya and matched the model’s location and body position to stills from the video footage. In Year 1, we used videos of live male *E. zonale* and *E. barrenense* as a reference to create a path for animations, which was a composite of 15s *E. barrenense* behavior and 15s *E. zonale* behavior. In Year 2, we used videos of live female *E. zonale* to create a path for animations, which was 47s of nose jabbing (typically performed prior to spawning) and swimming behavior. All animations in a given year followed the same path, i.e., displayed the same behavioral sequence. “Skins” were then applied over 3D models in Maya to provide conspecific or heterospecific features to models. Skins are 2D images that are wrapped around a 3D model so that the 2D image covers the 3D space of the model. After applying a path and skin to 3D models, animations were exported as a video file to be used in behavioral trials.

We created seven different types of skins for animations (Table 1). One was used to test baseline preference for conspecific animations; four different skin types were used to test the effect of focal region features on male mate choice; and, two were created as controls (details below). Skins were designed with *E. zonale* as the intended focal species, since previous studies have shown that this species responds to video and computer animations in a comparable manner to live stimuli (Roberts et al., 2017, 2019). To create conspecific skins, we randomly selected RAW photographs of female *E. zonale* from the Line Creek and Middle Fork Red River sites in Year 1 (N = 3 from each site) and from the Line Creek Site in Year 2 (N = 5). Heterospecific skins were created using RAW photographs of *E. barrenense* (N = 6 in Year 1; N = 5 in Year 2). RAW photographs were manipulated to generate the different skin types (Table 1) in Adobe Photoshop (Adobe Inc., San Jose, CA) prior to being applied to the 3D model in Maya. All skins were added to the female *E. barrenense* 3D model (with the exception of the baseline animations in Year1), so that the only difference between animations in a behavioral assay was the focal region feature.

**Table 1.**
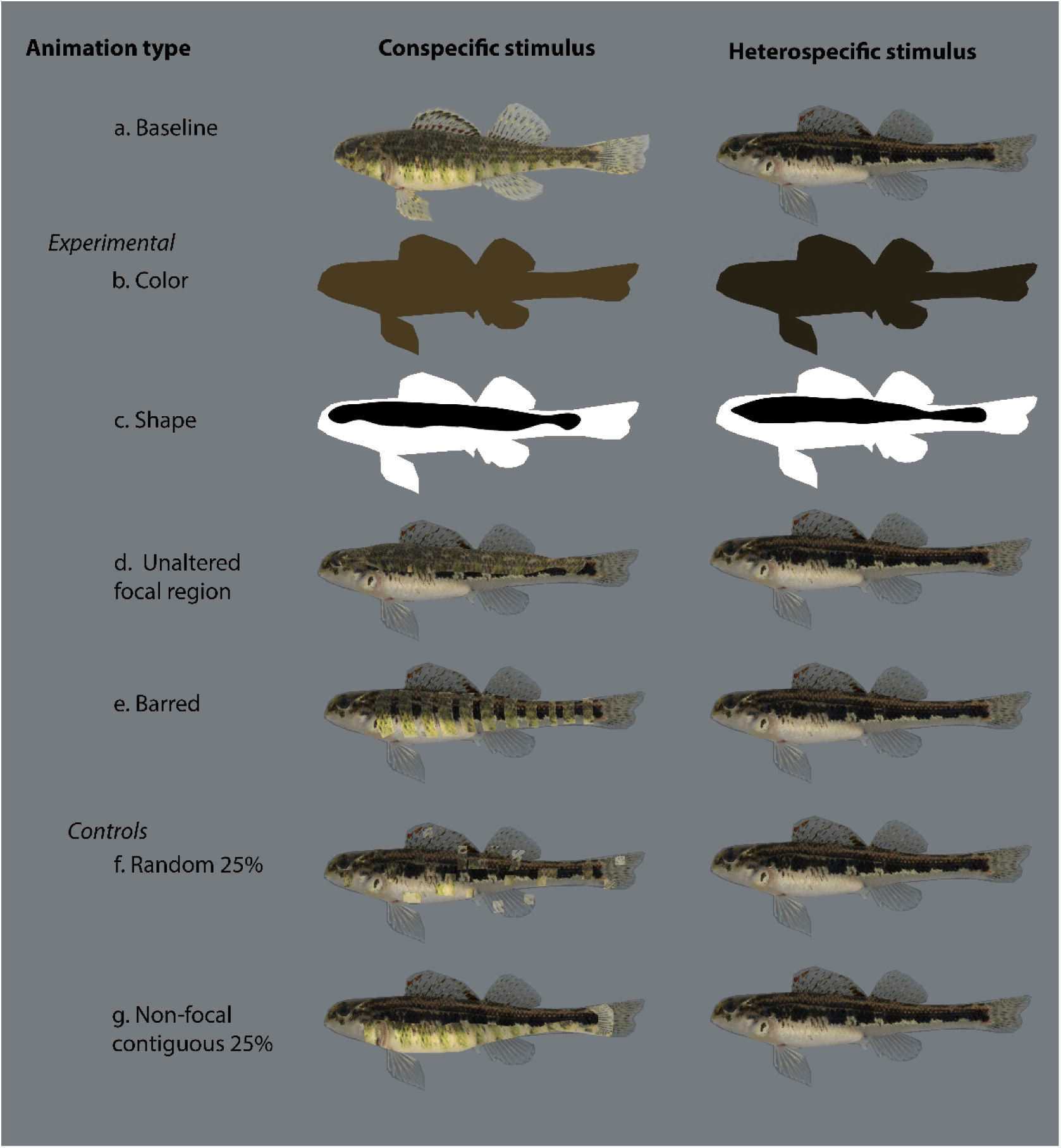
Examples of images used as skins to make 3D animated models for behavioral trials. (a) Unaltered conspecific female *E. zonale* and unaltered heterospecific female *E. barrenense* used in baseline trials. (b) Color-only models were created using the normalized average focal region color for conspecific and heterospecific coloration applied to 3D models. (c) Shape-only models used conspecific and heterospecific focal region shape in black on a white background applied to the female *E. barrenense* 3D model. (d) Full focal region model was created by applying conspecific *E. zonale* focal region over a heterospecific *E. barrenense* female image. (e) Barred model was created by applying the conspecific barred patterning over a heterospecific model. (f) Random area control was created by randomly selecting 100 x 100 pixel squares of conspecific *E. zonale* and applying these over a heterospecific image. (g) Non-focal area control was created by selecting a contiguous patch of conspecific *E. zonale* patterning excluding any area of the focal region and applying it over a heterospecific image. Stimuli consisting of a heterospecific base model with conspecific elements applied over top (d-g) were paired in behavioral trials with the unaltered heterospecific female.

### Baseline animations

To measure a male’s baseline preference for conspecific animations relative to heterospecific animations, we used unaltered photographs of female *E. zonale* and *E. barrenense* as skins (Table 1a). Measuring males’ baseline preferences provides a reference with which to compare male preference for the various conspecific features. We also used male response to baseline animations to select males to be used in the full study (see below). Since the goal of the current study was to determine which female features promote behavioral isolation, we only included males that showed strong conspecific preference in baseline trials.

#### Color-only animations

To create color-only animations, we measured the average RBG pixel value for female *E. zonale* and *E. barrenense* focal regions. The focal region RGB values were normalized so the sum of each independent color channel was equal between all photographs, allowing us to test the effect of focal region color independent of differences in overall brightness. The normalized average RGB values of conspecific and heterospecific focal regions were applied as the skins for animations. This resulted in animations that displayed either conspecific or heterospecific focal region color over the entire 3D model (Table 1b).

#### Shape-only animations

We created the shape-only animations by filling in the focal region area with pure black and applying pure white fill to the remainder of the skin (Table 1c). This was done for both conspecific and heterospecific focal region shape to generate conspecific shape-only animations and heterospecific shape-only animations. The shape-only skin was then applied to animations, allowing us to test the effect of focal region shape without any effect of color.

#### Unaltered focal region animations

To create the unaltered focal region animation, we took a photograph of a heterospecific female and pasted an *E. zonale* focal region over the original image (Table 1d). This skin was added to the models, resulting in an animation of a heterospecific female displaying a conspecific female focal region. The focal region animation allowed us to test whether the addition of the conspecific focal region to a heterospecific female increases male preference for an otherwise heterospecific animation.

#### Barred animations

Previous studies have shown that females prefer the barred patterning of males in *E. zonale* (Williams & Mendelson, 2011), and our pattern energy analysis indicated that the male barring pattern corresponds to the dominant marking size for males in this species (Fig. 3d). Since female *E. zonale* also display bars in muted green coloration (Fig. 1), we therefore created and tested an additional animation containing the barred patterning, which allowed us to test whether males and females prefer the same species-specific phenotype. We created the barred animation by manually selecting female *E. zonale* bars using the polygon selection tool in Photoshop and adding the barred patterning over a heterospecific female skin. This skin was applied to animations, resulting in animations of female *E. barrenense* displaying female *E. zonale* bars (Table 1e).

#### Random area control animations

The random area control was created by randomly selecting 100 x 100 pixel squares of female *E. zonale* patterning representing 25% of the total female body and applying these over an image of a female *E. barrenense* (Table 1f). Random area animations allowed us to control for any effect on male preferences of pasting conspecific elements over a heterospecific female, thus controlling for the skin editing process in the unaltered focal region and barred animations. The random area control also indicated whether conspecific pattern elements outside of those being directly tested in focal region and barred trials influence male preferences.

#### Non-focal area control animations

The non-focal area control was created by selecting a contiguous body area of a female *E. zonale*, excluding any area that was within the focal region, and applying this over a female *E. barrenense* skin. This resulted in animations that had 25% non-focal conspecific phenotype over an otherwise heterospecific female (Table 1g). Because focal regions were largely contiguous areas of female patterning, non-focal area animations allowed for a more direct control of the editing process on male preferences for the unaltered focal region animations. The non-focal area control also provides a reference for male preferences to 25% conspecific signal, to ensure that male preferences in focal region trials are not simply due to receiving sufficient conspecific signal, regardless of its location on the body.

### Behavioral assays

Experimental setup for the behavioral trials followed those used in Roberts et al. (2019), with a central test tank flanked on either short side by two computer monitors with two 10-cm “association zones” marked on each side of the tank closest to the computer monitors. Male *E. zonale* were placed in the test tank while the monitors displayed a grey screen for ten minutes to allow for acclimation to the testing environment. One monitor then began playing an animation of a conspecific female (or conspecific female feature) while the other monitor played an animation of a heterospecific female (or feature). Once the focal male had visited both 10-cm association zones and subsequently returned to the neutral zone (i.e. was not in either association zone), we began recording the amount of time a male spent in the conspecific and heterospecific association zones over a 10-minute trial period using JWatcher (Blumstein et al. 2000). From this, we calculated the proportion of time a male spent in the conspecific and heterospecific association zone relative to the entire trial time, and male strength of preference (SOP) after Stalker (1942). This is calculated as:

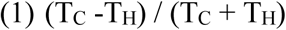

Where T_C_ is the absolute amount of time spent in the conspecific association zone and T_H_ is the absolute amount of time spent in the heterospecific association zone. SOP can range from +1 to - 1, with +1 indicating a complete preference for conspecific female animations (or features), −1 indicating a complete preference for heterospecific animations (or features), and zero indicating no preference for either type.

In Year 1, males were first subject to a baseline test in which they chose between an unaltered conspecific and heterospecific animation. These animations displayed the appropriate color-pattern and body shape of *E. zonale* and *E. barrenense* females. We then selected males that exhibited SOP > 0.15 to test the role of female focal features in male mate preference. Because there is variation in male preferences in darters (Mattson et al., in prep), selecting males with SOP above a threshold value helped ensure that we measured the effect of conspecific features on male preferences only in those males that exhibit a baseline conspecific preference. In Year 2, males were subjected to two baseline tests before selection. The first baseline test consisted of an unaltered conspecific and an unaltered heterospecific female. The second baseline test showed the same unaltered females as before, but with the side of the tank each female was presented on switched relative to the first baseline test. These animations displayed the appropriate color-pattern of conspecific and heterospecific females, but both stimuli displayed *E. barrenense* body shape. This helped to ensure that males not only preferred conspecific females, but were consistent in their preference across multiple trials, and that their preference was due to skin patterning rather than body shape. Applying an unaltered conspecific skin to the 3D *E. barrenense* model also provided a better comparison to male preferences for experimental animations, which were created using the female *E. barrenense* 3D model. To select males to be used in the full study in Year 2, we summed the amount of time spent in the conspecific and heterospecific association zones across both baseline trials. The summed time spent in the conspecific and heterospecific association zones was applied to equation 1. Males that had SOP > 0.15 were then selected to be used in further trials. In Year 1, we showed N = 18 males the baseline animation and N = 8 met the criteria for inclusion in the full study. In Year 2, we showed N = 25 males two sets of baseline animations and N = 12 met the criteria for inclusion in the full study.

In Year 1, we presented selected males with color-only animations, shape-only animations, an unaltered focal region animation, and a random area control animation, in random order. In Year 2, we showed selected males a barred animation, an unaltered focal region animation, and a non-focal area control animation in random order (Table 1). For the color-only and shape-only animations in Year 1, fish saw the conspecific feature on one screen and the heterospecific feature on the other screen. For the unaltered focal region, barred, and both control animations in Year 1 and Year 2, because they were created by adding conspecific features over a heterospecific skin, fish saw the conspecific feature on one screen and the unaltered heterospecific animation on the other. The side of the tank that the conspecific feature was presented on was switched between all trials. We then calculated the proportion of time spent in the left-hand and right-hand side association zones relative to the total amount of time spent in both association zones. Two males from Year 2 spent more than 70% of their total association time on one side of the tank across all trials (N = 20) and were removed from analysis for potential side bias.

For behavioral assays, we first assessed whether there was a significant effect of year on SOP for the full dataset using a linear mixed model, taking into account repeated individual ID as a random effect. For baseline and unaltered focal region trials, which were the only two treatments conducted in both years, we additionally tested for differences between years by comparing SOP using unpaired t tests or Mann-Whitney *U* tests for parametric and non-parametric data, respectively. To determine whether male *E. zonale* prefer conspecific focal region elements, we compared the proportion of time spent in the conspecific and heterospecific association zones for all trials. Association time has been found to be a proxy for mating preference in other species of fishes (Aspbury & Basolo 2002; Gonçalves & Oliveira 2003; Lehtonen & Lindstrom 2008; Jeswiet & Godin 2011) and association times in dichotomous choice assays are consistent with mating preferences in unrestricted stream trials for darters (Williams & Mendelson 2010; Martin & Mendelson 2013). We used a generalized least square (GLS) model to directly test for differences in SOP between the baseline preference trials, random area controls, and non-focal area controls. We also included in that model any experimental trials that showed a significant difference between conspecific and heterospecific association times in the dichotomous choice assays. The GLS model takes into account heterogeneous data and the repeated measure of ID across trial types. Post-hoc analyses to compare differences between groups were conducted using the lsmeans package in R with a Tukey correction applied for multiple comparisons. We also calculated the effect size using the effsize package in R for all post-hoc comparisons. We tested the assumption of normality for our data using Shapiro-Wilks and Q-Q plots and performed non-parametric statistics where appropriate. All statistical analyses were conducted in R.

## Results

### Quantifying focal regions: pattern energy analysis

Female *E. zonale* and *E. barrenense* had the highest pattern energy at the same filter size (440 pixels), indicating a similar dominant marking pattern. Pattern energy was higher for female E. barrenense than for female *E. zonale* at this filter size (t-test: t = 3.67, *P* = 0.0006; Fig. 2), indicating higher contrast for *E. barrenense* females in this dominant marking. Males of both species exhibited lower pattern energy overall than females (Fig. 2) and also differed from females in their dominant marking size: the filter size with maximum energy for male *E. zonale* was 172 pixels and for male *E. barrenense* was 275 pixels. Our analysis thus indicates that pattern is sex-specific, both in dominant marking size and in overall contrast, within each species.

### Behavioral assays

The results of the linear mixed model suggested no significant effect of year on SOP for the full dataset (t_16_ = - 0.33, *P* = 0.75). Further, comparing SOP in baseline and unaltered focal region animations revealed no significant difference in SOP between baseline trials between years (Mann-Whitney *U* test: *Z* = 1.15, *P* = 0.27) or between unaltered focal region trials between years (two tailed t-test: t = 0.61, *P* = 0.55). We therefore combined Year 1 and Year 2 data where possible for all further analyses.

Males spent significantly more time with the conspecific female than the heterospecific female in baseline trials, spending a mean ± SE of 49.07 ± 4.57% of total trial time in the conspecific association zone, compared to 23.00 ± 2.83% of total trial time in the heterospecific association zone (t = 3.87, *P* = 0.001). This significant difference is expected, since we selected males that preferred conspecific animations. We found no significant difference in the proportion of time spent in the conspecific or heterospecific preference zone for the random area control (t = −0.08, *P* = 0.94) and the non-focal area control (t = −1.36, *P* = 0.21). We also found no significant difference in the proportion of time spent in the conspecific or heterospecific association zone for the color-only (Wilcox signed-rank test: *Z* = −0.56, *P* = 0.64), shape-only (*Z* = −0.28, *P* = 0.84), or barred trials (paired t-test: t = −0.64, *P* = 0.54). Males spent significantly more time in the conspecific association zone for the unaltered focal region trials, spending a mean ± SE of 31.84 ± 4.48% of total trial time in the conspecific association zone, compared to 18.00 ± 2.78% of total trial time in the heterospecific association zone (t = 2.55, *P* = 0.02). Box and whisker plots for all comparisons are presented in Supplementary Figure 1. Unaltered focal region trials were therefore the only experimental animation type to be included in the GLS model.

Results of the GLS model showed a significant difference in SOP between animation types (F = 6.64, DF = 3, *P* = 0.0012). Male SOP was highest in the baseline trials (mean ± SE SOP of 0.42 ± 0.06), intermediate in the unaltered focal region trials (0.25 ± 0.12), and lowest in the random area (0.07 ± 0.24) and non-focal area controls (−0.20 ± 0.15). Post-hoc analyses showed that SOP in the unaltered focal region trials were not significantly different from either the baseline or the control trials after Tukey corrections (Fig. 4).

**Figure 4.**
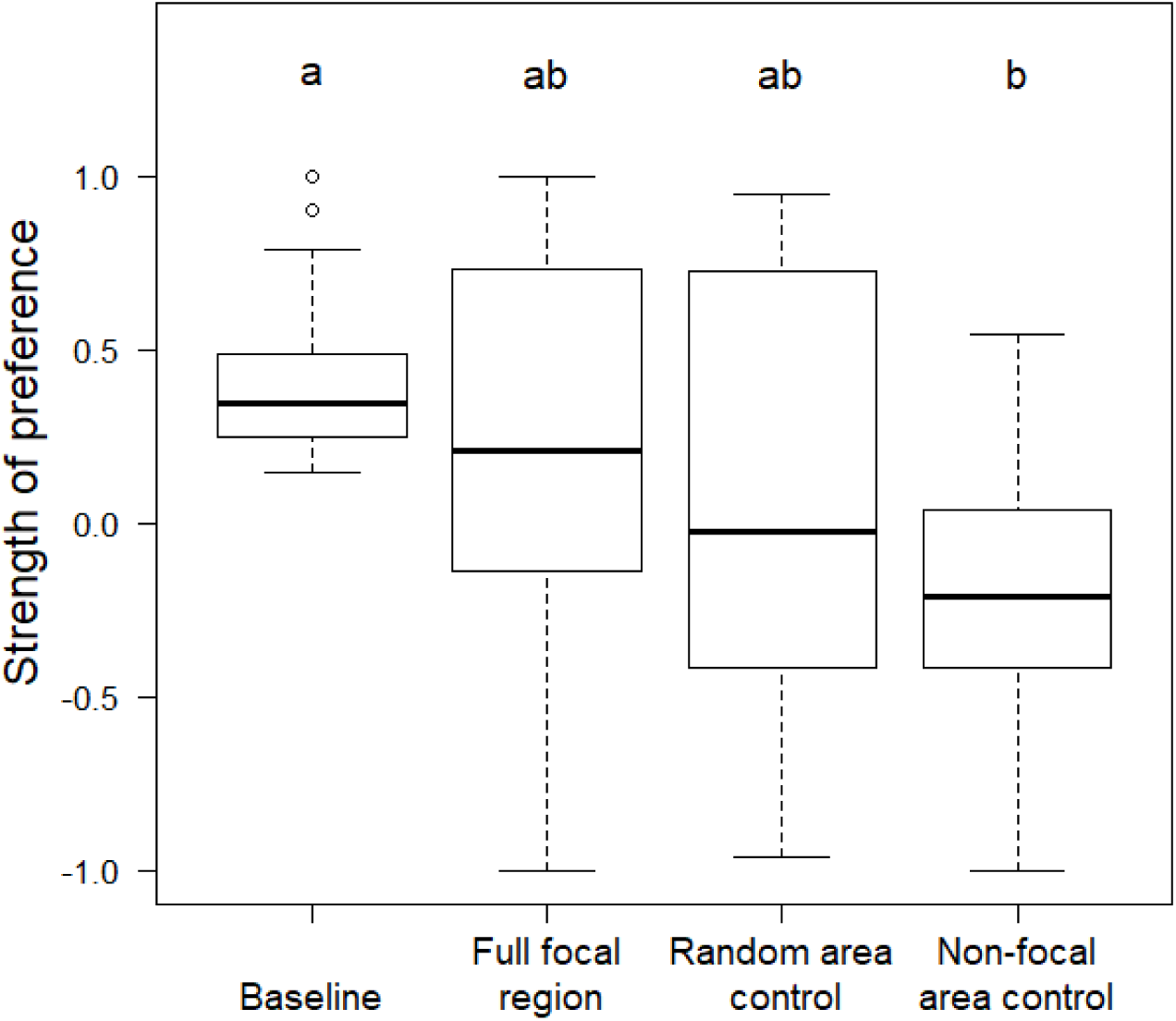
Strength of preference in baseline, control and unaltered focal region trials for male *E. zonale.* Positive strength of preference values indicate a preference for conspecific stimuli, negative values indicate a preference for heterospecific stimuli, and a score of zero indicates no preference. Bars represent medians, boxes indicate upper and lower quartiles, whiskers show sample minima and maxima, and open circles show outliers. Letters above boxes indicate a significant difference between groups at an alpha level of 0.05. Groups labeled with the same letter code are not significantly different from each other after applying a Tukey correction for multiple comparisons. Baseline and unaltered focal region trials from Year 1 and Year 2 were combined for N = 18; random area controls from Year 1 represent N = 8; and, non-focal area controls from Year 2 represent N = 12.

Considering effect sizes, we found a small effect (d = 0.36) of trial type comparing SOP between the baseline and unaltered focal region trials. There was a larger effect of trial type when comparing the two control trials to focal region trials (d = 0.49 (random area) and d = 0.69 (non-focal)), although these values both fall approximately into the intermediate effect level given the benchmarks suggested by Cohen (1988). There was also an intermediate effect (d = 0.68) of trial type when comparing the baseline to the random area control trials and, finally, a large effect when comparing the baseline to the non-focal area control trials (d = 1.17).

## Discussion

Our results suggest that male preference for conspecific females in *E. zonale* is mediated, at least in part, by a sex-specific patterning element that we identified as the female focal region. Males spent significantly more time with a heterospecific animation displaying the conspecific *E. zonale* focal region than with the same heterospecific animation without the conspecific focal region. We also found no significant difference between SOP for baseline animations and unaltered focal region animations (Fig. 4). While we also found no significant difference between SOP for the controls compared to the unaltered focal region trials (Fig. 4), there was a significant difference between baseline and the non-focal area control, suggesting that the unaltered focal region animation yielded an intermediate response. Notably, the effect size was larger for comparisons between the focal region and both controls than for the baseline-focal region comparison (1.4x and 1.9x larger), suggesting greater similarity in male response to unaltered conspecific females and focal region animations. Taken together, these data suggest that the focal region area may be an important female signal that affects male mate choice.

How female phenotypes influence male mate choice between species has been studied more extensively in insects, most notably in Lepidoptera. In *Heliconius cydno* and *H. pachinus*, males prefer females displaying conspecific coloration irrespective of pattern (Kronfrost et al., 2006). Males of the putative hybrid species *H. heurippa* prefer the red and yellow coloration of conspecific female coloration, with removal of either color significantly reducing male attraction towards female models (Mavárez et al., 2006). In these cases, traits that promote assortative mating are generally conspicuous traits and are present in both sexes. Female darters are, however, generally unornamented relative to males, making it unclear which female features might affect male preferences. The use of pattern energy analysis to objectively identify the dominant female pattern allowed us to test the role of female pattern elements on male mate choice. Notably, the female focal region identified by pattern energy analysis was different than the barred body patterning that functions in behavioral isolation via female mate choice (Williams & Mendelson, 2011), despite being shared between both males and females in this species. In fact, in trials where female barred patterning was added to heterospecific animations, there was no significant difference in male association time with the barred female and an unaltered heterospecific female, suggesting that this pattern element in females does not contribute to assortative mating by males.

Selecting the focal region following pattern energy analysis for photographs of male *E. zonale* did identify the barred pattern as the male focal region (Fig. 3), indicating that male and female *E. zonale* display sex-specific dominant patterning. In addition to functioning as a sexual signal in female mate choice, male barred patterning appears to be used as a signal in male-male aggression for *E. zonale*. Male *E. zonale* that were shown painted models representing conspecific and heterospecific male color-pattern (bars versus stripe, respectively) spent more time with conspecific models (Williams & Mendelson, 2013). Male association bias towards conspecific models is consistent with findings that male *E. zonale* more frequently chase conspecific than heterospecific males in artificial stream assays (Williams & Mendelson, 2010; Roberts & Mendelson, in review). Increased male aggression towards conspecific male signals might suggest that the female focal region differs from males to reduce sexual harassment by males. Female signals are thought to reduce male harassment in a number of species (Robertson, 1985; Galán, 2000; Godsen & Svennson, 2009; Hosken et al., 2016). For example, in a species of damselflies (*Ischnura elegans*), females exhibit heritable sex-limited color polymorphisms, with females of one morph displaying the vivid coloration present on males while the other female morphs are drably colored. Male mating harassment was shown to change in a density dependent manner based on the most common female color morph, suggesting that selective pressures to avoid male harassment can shape female signals (Godsen & Svennson, 2009).

The results of the discriminant function analysis suggested that focal region color and shape alone were both able to predict species identity significantly above chance, indicating that males could possibly use these signals as reliable indicators in mate choice. However, males did not prefer color-only or shape-only conspecific animations in our dichotomous choice assays. It is possible that color replication by the computer monitors were not good representations for actual female color, although a previous study in *E. zonale* found that computer monitors replicate the majority of darter body coloration similarly to live fish (Roberts et al., 2017). It may also be the case that males are not the intended receiver of these signals, and therefore would not be expected to discriminate between females based on those signals alone. For example, both male and female turquoise-brown motmots exhibit elaborate tail ornaments (*Eumomota superciliosa*). These elaborations appear to be a sexual signal in males, but there is no apparent function of elaborate female tails in male mate choice (Murphy, 2007b, 2008). Rather, evidence suggests that elaborate female tails function primarily as a deterrent against attack by predators (Murphy, 2006, 2007a). It is possible that female color and dominant pattern shape in *E. zonale* and *E. barrenense* are used in interactions with predators, in female-female interactions, or may be preferred by males when tested with a species other than the one tested in the current study. Alternatively, it is possible that these individual traits, although diagnostic of species using DFA, are not biologically relevant. Our results suggest that patterning within the focal region, rather than color or shape of the focal region, is necessary to elicit male preference and highlight the importance of using behavioral tests to verify biological function.

The design of dichotomous choice assays makes it impossible to distinguish between male attraction towards a stimulus and male aversion towards the alternative stimulus. Female focal regions identified for *E. zonale* were similar areas of the body to female *E. barrenense* focal regions (i.e. the flank above the lateral line). Thus, by creating focal regions animations in this study by pasting *E. zonale* focal regions over images of female *E. barrenense* added a conspecific signal to a heterospecific female, potentially making the animation more attractive to males relative to the unaltered heterospecific animation, but also masked a species-identifying heterospecific signal, potentially making the animation less aversive to males relative to the unaltered heterospecific animation. Regardless of whether male association time is mediated by attraction towards or repulsion from stimuli, association time in dichotomous choice assays are good predictors of spawning behavior (Aspbury & Basolo 2002; Gonçalves & Oliveira 2003; Lehtonen & Lindstrom 2008; Jeswiet & Godin 2011), including in *E. zonale* and *E. barrenense* (Williams & Mendelson, 2010; Roberts & Mendelson, in review). Males spent significantly more time with focal region animations than with unaltered heterospecific females in dichotomous choice assays, meaning that females with a conspecific focal region would likely experience higher reproductive success than heterospecific females. The use of computer animations in behavioral studies can allow future studies to determine if mating outcomes are determined by attraction or aversion towards conspecific and heterospecific signals.

In summary, we identified a region of female body patterning that appears to mediate behavioral isolation via male mate choice in a species of darter. These results highlight the importance of using objective measures of phenotype in studies of animal behavior. Male traits that are used in female mate choice tend to be those that are easily identified by a human observer in sexually dimorphic species. The female focal region identified for female *E. zonale* in the current study represents a body region that differs from the primary species-identifying patterning in male *E. zonale* (i.e. bars). These results illustrate that the evolution of female signals may be overlooked in sexually dimorphic species due to the misidentification of relevant female signals or because female-specific signals are simply assumed to be absent. Identifying female traits that are biologically relevant in male mate choice is a critical first step in understanding female trait evolution.

## Acknowledgements

We would like to thank S. Hulse for assistance in the field and with photography, T. Bhardvay and S. Parsa for their assistance in behavioral trials, and T. Cronin, R. Fuller, J. Leips, and K. Omland for their feedback on manuscript preparation. This research was funded in part by an Animal Behavior Society Student Research Grant provided to NSR.

## Author Contributions

NSR and TCM conceived and designed the experiment and collected field collections. NSR collected data and performed statistical analyses. The manuscript was written by NSR with input from TCM. Both authors gave approval of the final manuscript prior to submission.

---

**Supplemental Figure 1.**
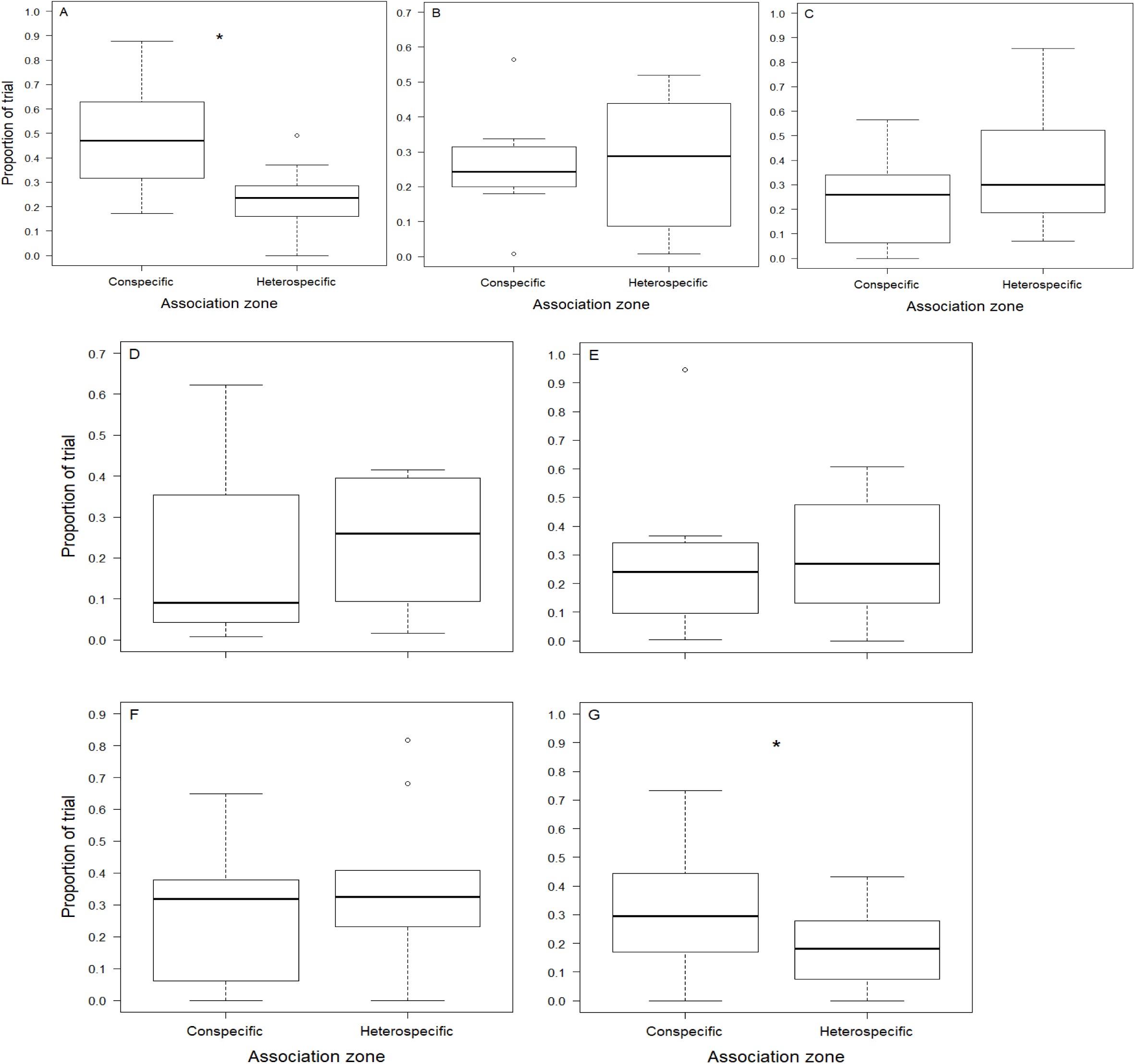
Proportion of time spent in the conspecific and heterospecific association zones for male *E. zonale*. (A) Males spent significantly more time in the conspecific association zone than in the heterospecific association zone in baseline trials. This was expected since we selected males with conspecific preference to be used in the full set of trials. There was no significant difference in the proportion of time spent with the conspecific animation compared to the heterospecific animation in (B) random area control trials, (C) non-focal area control trials, (D) color-only trials, (E) shape-only trials, or (F) barred trials. (G) Males spent significantly more time in the conspecific association zone relative to the heterospecific association zone in the unaltered focal region trials. Bars represent medians, boxes indicate upper and lower quartiles, whiskers show sample minima and maxima, and open circles show outliers. * Indicates a significant difference between groups.

## Notes

### Competing Interest Statement

The authors have declared no competing interest.

